# Respiratory depression and analgesia by opioid drugs in freely-behaving larval zebrafish

**DOI:** 10.1101/2020.09.30.320267

**Authors:** Shenhab Zaig, Carolina Scarpellini, Gaspard Montandon

**Affiliations:** Keenan Research Centre for Biomedical Sciences. St. Michael’s Hospital. Unity Health Toronto; Department of Medicine, Faculty of Medicine, University of Toronto

## Abstract

An opioid epidemic is spreading in North America with millions of opioid overdoses annually. Opioid drugs, like fentanyl, target the mu opioid receptor system and induce potentially lethal respiratory depression. The challenge in opioid research is to find a safe pain therapy with analgesic properties but no respiratory depression. Current discoveries are limited by lack of amenable animal models to screen candidate drugs. Zebrafish (*Danio rerio*) is an emerging animal model with high reproduction and fast development, which shares remarkable similarity in their physiology and genome to mammals. However, it is unknown whether zebrafish possesses similar opioid system, respiratory and analgesic responses to opioids than mammals. In freely-behaving larval zebrafish, fentanyl depresses the rate of respiratory mandible movements and induces analgesia, effects reversed by mu-opioid receptor antagonists. Zebrafish presents evolutionary conserved mechanisms of action of opioid drugs, also found in mammals, and constitute amenable models for phenotype-based drug discovery.

## Introduction

The current North American opioid epidemic is staggering, with millions of emergency visits and over 45,000 deaths annually (Florence, Zhou, Luo, & Xu, 2016). Synthetic opioids like fentanyl and oxycodone, or natural opioids like heroin and morphine are highly addictive (Wilkerson, Kim, Windsor, & Mareiniss, 2016) and can lead to respiratory depression (Dahan, Aarts, & Smith, 2010; (Montandon, Cushing, et al., 2016; (Nagappa, 2017), that can be lethal with overdose (Gomes & Juurlink, 2016; (Jones, Mack, & Paulozzi, 2013). Opioids are critical pain therapies and there are currently no other medications that offer effective relief of pain (IASP, 2018). However, the side-effects of opioids limit their effective use and may lead to sub-optimal pain treatments.

Opioid medications act on µ-, δ-, and κ-opioid receptors (Gutstein, 2001), which are expressed in discrete brain circuits. Although a wide range of opioids targeting these opioid receptors exist, the most potent opioids, such as fentanyl and oxycodone, are largely selective for the µ-opioid receptors (MORs) (Sora, Funada, & Uhl, 1997). Because MORs are expressed in neural circuits involved in breathing and nociception, opioids present effects beyond their intended therapeutic analgesic properties. MOR drugs induce side-effects such as addiction (Le Merrer, Becker, Befort, & Kieffer, 2009), sedation (Montandon, Cushing, et al., 2016), and hypoventilation (Dahan et al., 2010; (Montandon & Slutsky, 2019; (Nagappa, 2017). Hypoventilation is characterized by reduced respiratory rate (Macintyre, Loadsman, & Scott, 2011), reduced airflow (Ferguson & Drummond, 2006), prolonged apneas (Mogri, Khan, Grant, & Mador, 2008; (Nagappa, 2017), and severe hypoxemia (Brown, Laferriere, Lakheeram, & Moss, 2006). Complete respiratory arrest and bradycardia are the main causes of death by opioid overdose (Pattinson, 2008). Importantly, respiratory depression and cessation of breathing with opioid overdose are due to the action of opioid drugs on brainstem respiratory network (Montandon & Slutsky, 2019). The only available treatment to reverse respiratory depression during an opioid overdose is the MOR antagonist naloxone which directly blocks the binding of opioid ligands to MORs. However, naloxone also blocks the analgesic properties of opioid drugs, so it cannot be used as an adjunct treatment. The discovery of novel opioid drugs or combinations of drugs with potent analgesic properties without the side-effects of respiratory depression has been hindered due to the lack of animal models allowing behavioral assessments of pain and breathing, while allowing for large-scale drug discovery.

Identification of new molecular targets and drugs is critical to the development of safe opioid therapies. Due to the complexity of the machinery regulating MOR inhibition, wide scale screening of drugs acting on the known mechanisms regulating MOR inhibition in rodent models is not feasible. It is therefore critical to identify amenable model systems where large-scale screening can be developed, while preserving the complexity of vertebrate central nervous system and behaviours. The challenge is to quantify complex phenotypes like respiratory neural activity and nociception on a large scale, which is normally limited by the complexity of the animal models. Rodents are not optimal because their large size, generation time, and drug availability limit screen scale. An alternative is the larval zebrafish, which shares anatomy, physiology, and large parts of the genome with humans. The small size of larval zebrafish, ease of production, and their external fertilization which allows easy gene knockdown, makes the zebrafish an attractive model organism to screen genes and drugs. Here, we propose to use the larval zebrafish as simple, yet powerful, animal model, so that the molecular mechanisms of MOR respiratory and nociceptive inhibition can be identified. Importantly, the zebrafish larvae can also be used for large-scale drug screening to identify new drug candidates.

The larval zebrafish has emerged as an ideal model system for drug and gene discovery since it combines the biological complexity of *in vivo* models with a nervous system homolog to humans including a respiratory neural network. Although fish use a different strategy to absorb oxygen and eliminate carbon dioxide than mammals, they rhythmically produce mandibular movements to move water through their gills. In lampreys, the paratrigeminal respiratory group (pTRG) generates rhythmic mandible movements (Bongianni, Mutolo, Cinelli, & Pantaleo, 2016). Because of their close evolutionary origins (Missaghi, Le Gal, Gray, & Dubuc, 2016), the pTRG shares similarities with the mammalian respiratory network (Cinelli et al., 2013) which generates breathing and regulates respiratory depression by opioids (Montandon et al., 2011). The pTRG presents similar functional properties to that of mammals such as sensitivity to substance P (Mutolo, Bongianni, Cinelli, & Pantaleo, 2010). Here, we propose that respiratory mandible movements can be used as an index of respiratory network activity, similar to respiratory activity of the trigeminal muscle in mammals (Jacquin, Sadoc, Borday, & Champagnat, 1999). The zebrafish also possesses a nociceptive system encompassing spinal cord, brainstem and sub-cortical circuits (Taylor et al., 2017), with homology to mammals. Although pain circuit activity and its response to opioids cannot be directly assessed, the swimming escape response to nociceptive stimuli can be easily assessed in larval zebrafish. Swimming response to chemicals, such as formalin (Magalhaes et al., 2017) or acetic acid (Lopez-Luna, Al-Jubouri, Al-Nuaimy, & Sneddon, 2017) can be measured, as well as response to thermal stimuli (Malafoglia et al., 2014). Here, we propose to determine whether respiratory depression and analgesia can be measured in larval zebrafish.

Most opioid analgesics and drugs of abuse elicit their effects by binding to MORs (Gutstein, 2001). In zebrafish, the MOR has high homology to the mammalian MOR and shares 74% of its amino acid sequence with its mammalian counterpart (Sanchez-Simon & Rodriguez, 2008). The MOR also possesses similar binding properties to the mammalian MOR for morphine and DAMGO (Marron Fdez de Velasco, Law, & Rodriguez, 2009; (Mutolo, Bongianni, Einum, Dubuc, & Pantaleo, 2007). Because of the homologies of the respiratory, pain, and opioid systems in zebrafish and mammals, we propose that the larval zebrafish may mimic respiratory depression and analgesia by opioids and several other phenotypes associated with opioid drugs in mammals. The objectives of this study are to demonstrate that larval zebrafish have evolutionary conserved opioid properties mimicking mammals and that they can be used to investigate opioid-induced respiratory depression and analgesia. We aim to demonstrate that larval zebrafish replicates the opioid pharmacology observed in mammals and humans and that current pharmacotherapies to block respiratory depression can be tested in larval zebrafish.

## Methods

### Animal husbandry

Animal practices and experiments in adult fish, larvae, and breeding pairs followed laboratory standards (Avdesh et al., 2012) and were carried out according to the procedures outlined by the Canadian Council on Animal Care and were approved by St. Michael’s Hospital animal care committee. Only wildtype AB strain zebrafish larvae at 12-14 days post-fertilization (dpf) were used for experiments, except when comparisons between strains were made. Tübingen (TU) and crosses between TU and AB were used to compare strains. All fish were housed on a 14/10-hour light/dark cycle and kept at a constant water temperature of 28°C ± 0.5°. Larvae and adults were originally obtained from the Hospital for Sick Children and the University of Toronto Mississauga (Toronto, ON). For breeding, male and female adult fish at 4 months of age were placed in a breeding tank and separated by a divider. The next day the divider was removed and fish mated within the first half hour of the lights-on period and were returned to the rack (Aquaneering, CA, United States). The eggs were collected and placed in petri-dish filled with E2 embryo medium (NaCl, 15.0 mM; KCl, 0.5 mM; MgSO_4_, 1.0 mM; CaCl_2_, 1.0 mM; Na_2_HPO_4_, 0.05 mM; KH_2_PO_4_, 0.15 mM; NaHCO_3_, 0.7 mM). Unfertilized eggs were removed.

Beginning at 5-days post-fertilization, larvae were fed with Ziegler AP100 (artificial plankton) dry larval diet (100 microns) and were moved to 0.8 litre tanks filled with system water, with a density of 20 fish/100mL and were kept there until the experiment. Water quality was kept at a pH of 7.5±0.5 and with a conductivity of 500-1000ppm. Dissolved oxygen was maintained at 6-7ppm. Nitrites and ammonia were kept at <150ppm. All experiments were undertaken during daylight hours and fish were placed in an incubator at 28.5±0.5°C.

### Drug treatments

All opioid drugs were used with Health Canada approval. Fentanyl citrate and morphine HCl were obtained from Sandoz (QC, Canada). Naloxone hydrochloride was obtained from Mayer. The MOR antagonist CTAP (D-Phe-Cys-Tyr-D-Trp-Arg-Thr-Pen-Thr-NH_2_, lidocaine, the Ampakine CX614, and the 5-HT_4A_ receptor agonist BIMU8 were obtained from Tocris (ON, Canada).

### Determination of mandibular respiratory movements

Respiratory mandible movements were quantified in live zebrafish larvae using a custom-made system including a 4K high-definition camera (Basler 4K, 4096pi x 3000pi, model acA4112-30um, Edmunds Optics) and partially telecentric 7x zoom lens (Edmund Optics). The water was kept at a constant 28.5°C temperature using a heating pad. With this system, simultaneous recordings of 12 zebrafish larvae can be performed. As an index of respiratory network activity, mandible movements were measured and respiratory rate was quantified (**Figure 1a**). In a custom-made 12-well clear plate, larval zebrafish were placed in wells (diameter 10mm, depth 4mm) containing 60 µL of embryo medium. Using our video recording system, a dorsal video was performed for each drug combination. We used two approaches to determine the rate of mandible movement. Rate of mandible movements was visually counted by two independent researchers, blind to the drug combinations. Using this approach, the number of mandible movements per minute was calculated for each condition and each animal. In a subset of zebrafish larvae, mandible movements were also quantified by measuring pixel intensity changes in a defined area around the fish head. Tracings of pixel intensity over time were plotted and the rate of intensity changes was quantified using a custom-made software in Matlab (Mathworks, US). The validity of visual quantification was then compared to software quantification.

**Figure 1.**
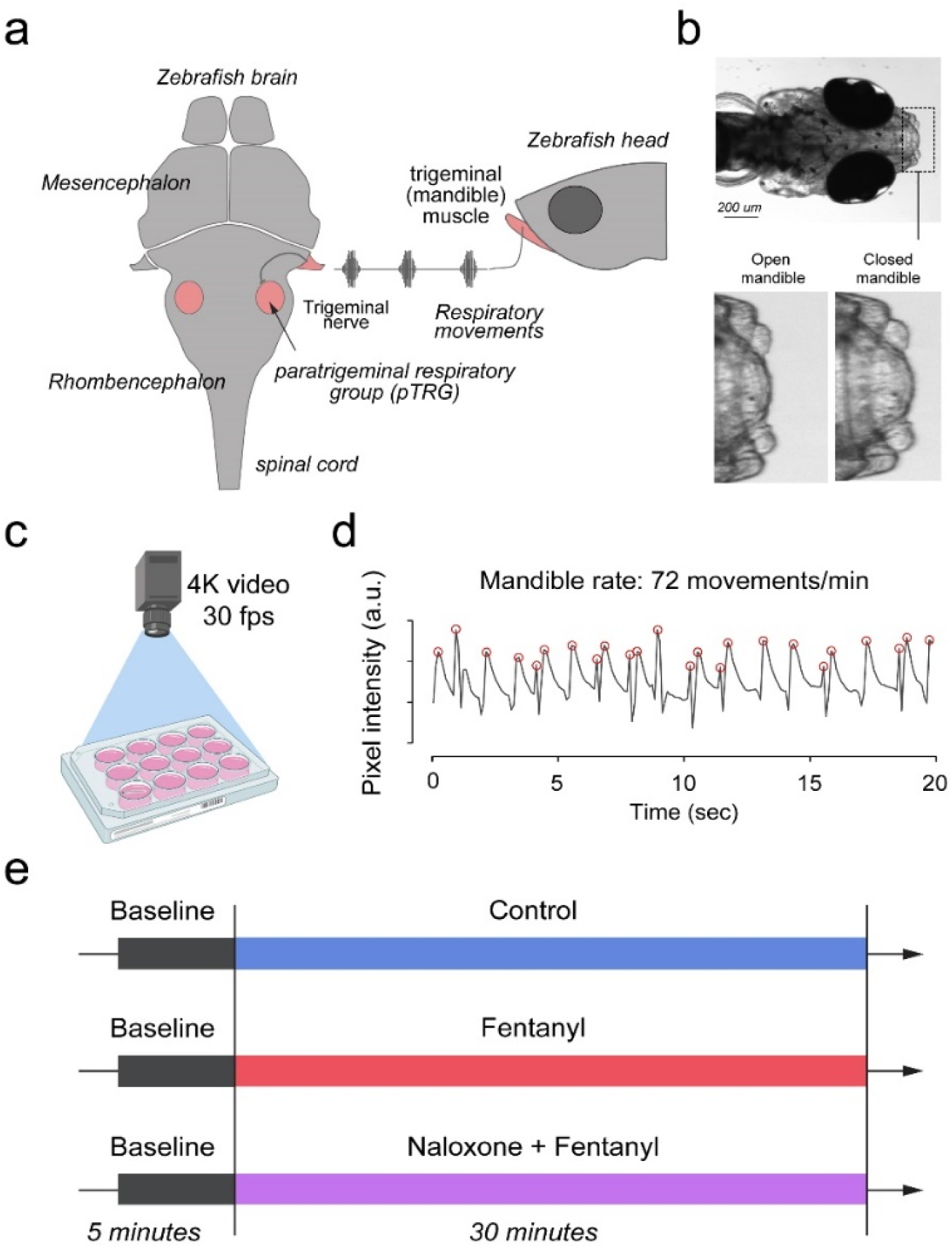
Respiratory network activity quantified using mandible movements in freely-moving larval zebrafish. The brainstem respiratory network of the larval zebrafish produces respiratory mandible movements (**a**). Respiratory mandible movements are used as an index of respiratory network activity (**b**). Mandible movements were quantified by looking at the pixel changes in the region of the mandible using a 4K high-definition camera (**c**). Respiratory rate was quantified in 12-14 day post-fertilization larvae positioned in 12-well plates (**d**). Red circles indicate peaks of mandible movements. To quantify respiratory rate depression induced by opioid drugs, fish were exposed to a drug treatment containing embryo medium only (control), fentanyl, or a combination of naloxone and fentanyl (**e**).

### Respiratory depression by opioids in zebrafish

Zebrafish larvae were positioned in wells and were left for 10 minutes to acclimatize to the environment. A video of baseline respiratory movement was then taken for 1 minute. Following the baseline recordings, combinations of drugs were applied to each well in a volume of 6 µL per well. Drugs were allowed to incubate for 5 minutes and a video was recorded for 30 minutes. To determine whether various drugs induce respiratory rate depression, we compared the rates of mandible movements between several groups of larvae: control group (embryo medium), fentanyl group, fentanyl/naloxone group (combination of fentanyl and naloxone), fentanyl/other drugs (fentanyl and potential drug candidates). All these groups were tested simultaneously in 12-well plates (4 larvae per group if 3 groups were tested). By combining all groups in the same multi-well plate, we ensured that all animals experienced the same conditions therefore allowing comparisons between groups. Respiratory rate values collected in larval zebrafish were normalized according to baseline mandible rates acquired before drugs were administered. Normalization considerably reduced the high variability encountered in larval zebrafish. Larval zebrafish may present different growth and behaviors at 12-14dpf due to variable development rates and access to food and nutrients within the tank. To consistently select larvae at similar development stages, we established selection criteria. For respiratory assays, larvae with rates of mandible movements below 40 movements/min at baseline were not considered because their respiratory network may not be properly developed.

### Swimming behaviour quantification

Nociception was assessed by measuring the swimming escape response to nociceptive stimuli. Zebrafish were placed individually into wells of a custom-made 24-well plate containing 90µL of embryo water. The well plate was made of white acrylic with a back-light to allow video recording. The plate was heated at 28.5°C. The same custom-made video recording apparatus, as mentioned above, was used to record swimming in larval zebrafish. Swimming experiments were analyzed using a commercial software (EthoVision XT v11, Noldus Information Technology, Netherlands), and swimming velocity (mm/sec) and angular velocity (degrees/sec) were quantified. Before applying drugs to the plate wells containing larvae, we measured a 10 min baseline. Larvae that swam less than 100 mm over the 10 minute baseline period (<0.17 mm/sec) were not considered for analysis. Formalin was then used as nociceptive stimulus. Such stimulus was previously validated in mammalian and adult zebrafish (Magalhaes et al., 2017) studies. We applied formalin in combination with opioid analgesics to determine opioid analgesia in larval zebrafish.

### Opioid analgesia in larval zebrafish

Zebrafish larvae (12-14dpf) were positioned in wells of a 24-well plate and left 10 minutes to acclimatize. A video of the baseline movements was taken for 10 minutes. Following baseline recordings, combinations of drugs were applied to each well in a volume of 10 µL and video was recorded for 20 minutes. To determine whether various drug combinations induce analgesia, we compared the swimming velocity between several groups of larvae: control group (embryo medium was administered), formalin group, fentanyl/formalin group (combination of fentanyl and formalin), formalin/fentanyl/naloxone. All these groups were tested simultaneously in 24-well plates (6 larvae per group if 4 groups were tested). The velocity calculated for the last 3 minutes of baseline (minutes 7-10) and first 3 minutes of treatment (minutes 0-3) were used for analysis.

### Statistical analysis

Data in larval zebrafish are often not normally distributed. Therefore, normality was tested using a Shapiro-Wilk method and equality of variances using the Brown-Forsythe test. When data distribution was not normal, we compared experimental groups using Wilcoxon rank tests and All Pairwise Multiple Comparison Procedures (Dunn’s method) as *post-hoc* tests. For respiratory rate, for instance, we used non-parametric tests and presented data as individual data points and medians with error bars showing 25^th^ and 75^th^ percentiles (interquartile range). When data was normalized according to baseline measurements, data followed a normal distribution. We then compared multiple groups using a one-way ANOVA followed by Holm-Sidak *post-hoc* tests to compare individual groups. For normal data, we presented data with means and standard deviation as error bars. In some cases, data presented clear outliers. To objectively identify outliers, we selected all data points below and above 1.5 x the interquartile range and eliminated these data points from the data set (Michel, Murphy, & Motulsky, 2020). All statistical tests and graphs were done using Sigmaplot (version 14, SAS) and figures were prepared with Adobe Illustrator (Creative Suite 5, Adobe).

## Results

Zebrafish use gills to promote gas exchange with water. Flow of water through the gills is generated by a complex respiratory network in the brainstem which presents similar anatomy and properties to the mammalian respiratory network (**Figure 1a**). The paratrigeminal respiratory group generates rhythmic movements of muscles involved in branchial movements. One of these muscles, the trigeminal muscle controls opening and closing of the mandible. Mandible movements can be easily recorded using a high-definition camera positioned on top of the swimming zebrafish (**Figure 1b, c**), and may be a robust index of respiratory network activity. Mandible movements can be quantified by counting the changes in pixel intensity around the mandible area (**Figure 1b, c**) and can be displayed over time (**Figure 1d**). The rate of mandible movements, identified as the number of mandible movement peaks (red circles, **Figure 1d**), indicates respiratory network rhythm and is considered here as an index of respiratory rate. Using the convenience of multi-well plates, we administered combinations of drugs to embryo medium as described in **Figure 1e**. Combinations of drugs were administered to separate groups of fish. Each animal only received one combination of drugs and did not receive consecutive combinations of drugs because it would not be feasible to remove drugs from the water.

To determine whether zebrafish can be used as a model of opioid-induced respiratory rate depression, we administered embryo medium (control) or fentanyl - a commonly used opioid analgesic-to larval zebrafish while quantifying the rate of mandible movements (**Figure 2a**). Larval zebrafish presented rates of mandible movements ranging from 15 to 115 movements per minute. Larvae were exposed to a concentration of fentanyl of 1µM and respiratory rate was recorded over a 30-min time period (**Figure 2b**). In the control group, respiratory rate initially increased compared to baseline but did not significantly change during the 30 minutes following baseline (**Figure 2b**). In the fentanyl group, respiratory rate was strongly reduced 4 minutes after fentanyl administration. Since respiratory rate depression was observed 5 minutes, we compared respiratory rates (measured over a one-minute time period between minutes 5 and 6) of groups exposed to embryo medium (control) and different concentrations of fentanyl (**Figure 2c**). The rates of mandible movements were not normally distributed (Shapiro-Wilk test: *P<0*.*05*, n>10 for all groups, **Figure 2c**). Fentanyl (0.2, 0.4 µM) did not significantly decrease rate of mandible movements compared to control (*P=0*.*207* and *P=0*.*099*, Kruskal-Wallis one-way ANOVA on ranks, **Figure 2c**). Fentanyl (1 and 3 µM) significantly decreased rate compared to control groups (*P=0*.*049* and *P=0*.*007*, **Figure 2c**). When data were normalized according to baseline rate, variability amongst groups was significantly reduced (**Figure 2d**), and data followed a normal distribution. Fentanyl significantly decreased respiratory rate at 0.2 µM (*P=0*.*049*), 0.4 µM *(P=0*.*042*), 1 µM (*P=0*.*010*) and 3 µM (*P<0*.*001*) compared to the control group (**Figure 2e**). In summary, fentanyl induced respiratory rate depression was observed at all concentrations of fentanyl tested when data was normalized according to baseline rates. We determined that such normalization best represented respiratory rate depression by fentanyl and we only presented normalized data for figures with respiratory depression.

**Figure 2.**
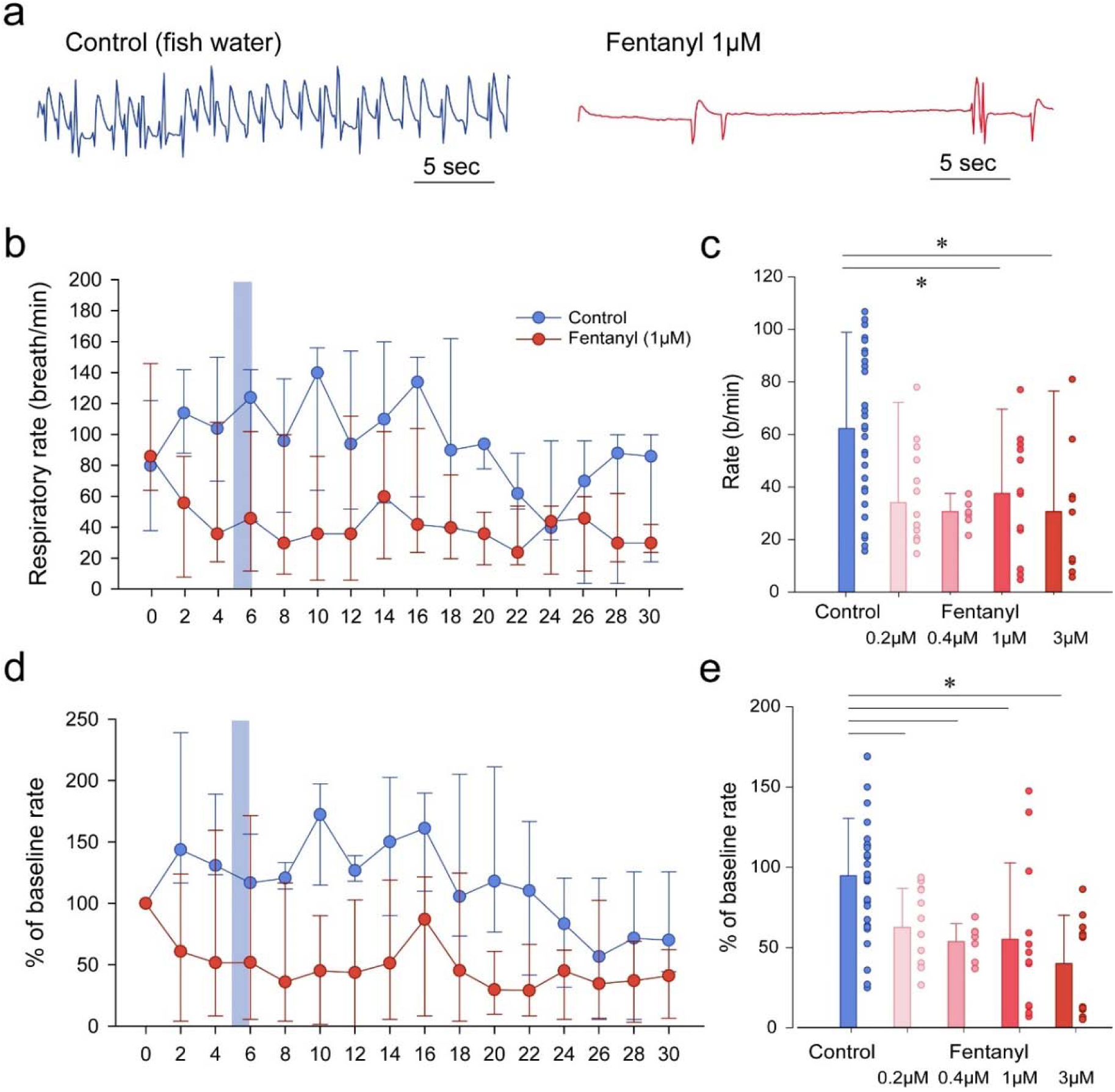
Respiratory rate depression by the opioid analgesic fentanyl in larval zebrafish. Representative mandible movements in larvae exposed to embryo medium or a solution of fentanyl (1µM). The rate of mandible movements was significantly decreased by fentanyl (**a**). Larvae were exposed to fentanyl over a period of 30 minutes. Significant respiratory rate depression was observed within 4 minutes following fentanyl application (**b**). Respiratory rate data did not follow a normal distribution and data are presented as medians with bars representing interquartile range. Increasing the concentration of fentanyl induced dose-dependent decreases in respiratory rate (**c**). Fentanyl produced significant decreases in respiratory rate at 1 µM (n=12) and 3 µM (n=11), but not at 0.2 µM (n=12) and 0.4 µM (n=7) compared to controls (n=32) **(c)**. Rates of mandible movements were not normally distributed due to the high variability in respiratory rates. To present more homogenous data, respiratory rates were normalized according to the baseline rate measured before drugs were applied to better represent how drug exposure changed rate **(d)**. Shading indicates time periods used to calculate average data in pan c & e. Fentanyl produced dose-dependent respiratory rate decreases at all concentrations tested. Normalized data are presented as medians ± 25^th^ and 75^th^ percentiles. Circles indicate individual data points for each zebrafish measured. * indicates significantly different medians compared to control with *P<0*.*05*.

To determine the pharmacology related to the MOR, we compared groups of fish exposed to fentanyl with fish exposed to fentanyl and the MOR antagonists naloxone and CTAP (**Figure 3**). Exposure to fentanyl reduced the rate of mandible movements (**Figure 3a**). The MOR antagonist naloxone at 5 µM did not significantly reverse respiratory rate depression induced by fentanyl (*P=0*.*237*), but showed a trend toward reversing fentanyl depression. The selective MOR antagonist CTAP significantly reversed respiratory rate depression by fentanyl (*P=0*.*049*, **Figure 3b**). A higher concentration of naloxone (20 µM) in combination with fentanyl (1 µM) showed increased respiratory rate compared to fentanyl alone (*P<0*.*001*, **Figure 3c**). In summary, fentanyl induced a significant respiratory rate depression which was blocked by the MOR specific antagonist CTAP and naloxone at 20 µM.

**Figure 3.**
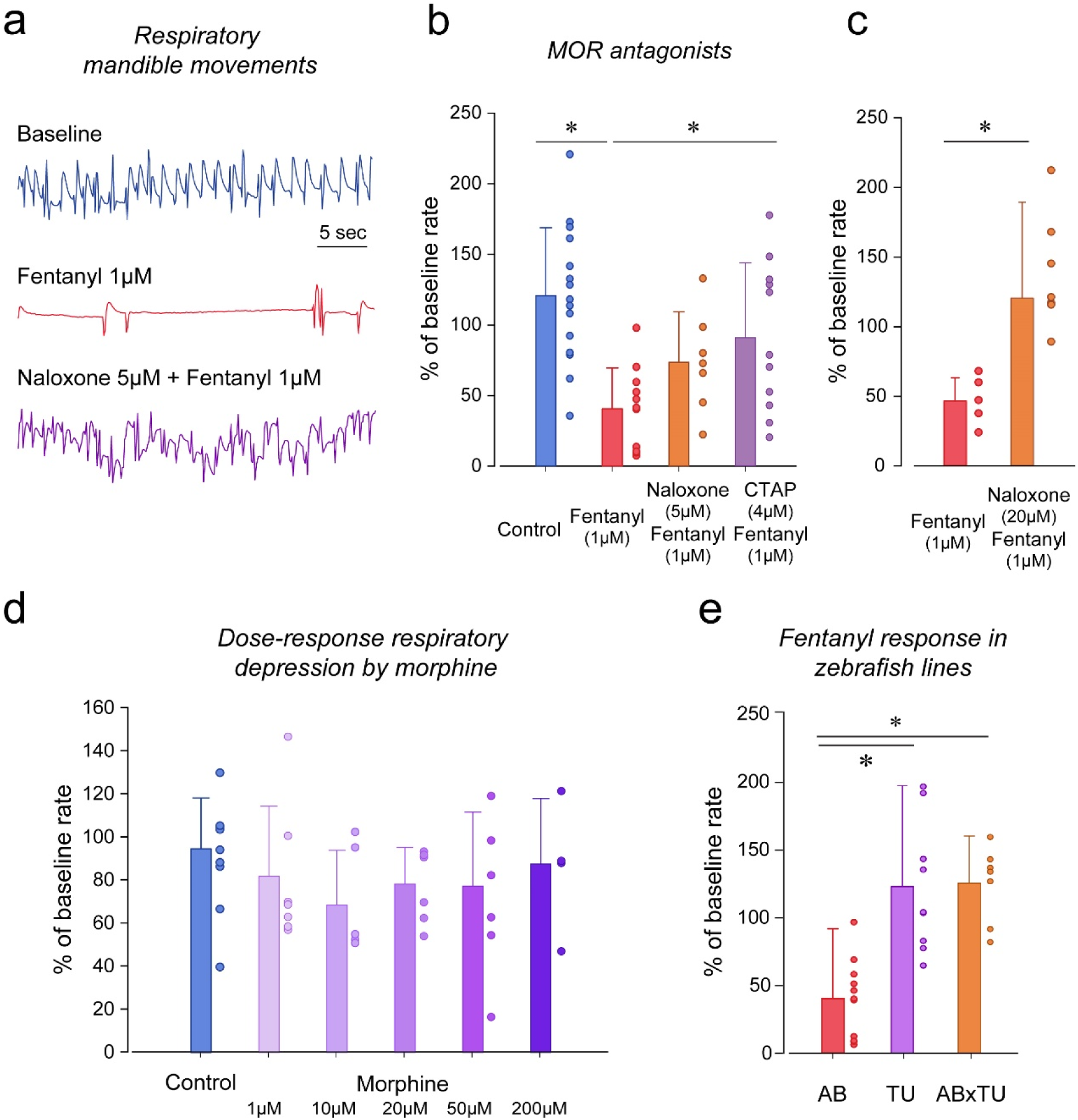
Opioid receptor pharmacology regulating respiratory depression in larval zebrafish. Representative tracings of mandible movements in larval zebrafish showing fentanyl (1 µM) reducing respiratory rate and reversal by the MOR antagonist naloxone (5 µM) (**a**). Fentanyl (1 µM) significantly decreased rate of mandible movements (n=11) compared to the control group (n=15), an effect significantly reversed by the highly selective MOR antagonist CTAP (4 µM, n=11) and with a trend toward significance for naloxone at 5 µM (n=10) (**b**). A higher concentration of naloxone administered with fentanyl (1 µM) blocked respiratory rate depression (n=6) when compared to fentanyl alone (n=5) (**c**). Morphine administered at various concentrations did not decrease respiratory rate compared to controls (**d**). Respiratory rate depression due to fentanyl was observed in the AB strain (n=11), but not in TU (n=9) or crosses between AB x TU (n=7) (**d**). * indicates significantly different medians compared to control with *P<0*.*05*. All data are presented as means with error bars showing ± standard deviation. Circles indicate individual data points for each zebrafish measured. * indicates groups significantly different. AB, AB zebrafish line. TU, Tübingen zebrafish line.

Morphine is a widely used opioid drug that induces less severe respiratory depression than highly potent opioids such as fentanyl (Gutstein, 2001). We tested whether morphine induces respiratory rate depression in zebrafish. Morphine at 1, 10, 20, 50 and 200 µM did not significantly depress respiratory rate when compared to the control group (*P=0*.*407*, **Figure 3d**). Because zebrafish strains may present genetic variability and different sensitivities to opioid drugs (Marron Fdez de Velasco et al., 2009; (Sanchez-Simon & Rodriguez, 2008), we compared respiratory depression by fentanyl in various zebrafish lines (**Figure 3e**). Fentanyl (1 µM) significantly reduced respiratory rate in AB fish (*P<0*.*001*), but not in TU (*P=0*.*770*) or ABxTU zebrafish crosses (*P=0*.*821*). To conclude, only the AB zebrafish strain presented sensitivity to fentanyl and was used for subsequent experiments.

To determine whether breathing in zebrafish larvae can be stimulated by specific excitatory drugs as observed in mammals, we administered two respiratory stimulants to zebrafish previously shown to stimulate breathing when depressed by opioids. We first administered the 5-HT_4A_ serotoninergic ligand BIMU8 (Manzke et al., 2003) with or without fentanyl (**Figure 4a**). BIMU8 combined with fentanyl did not show reversal of respiratory rate when compared to the fentanyl group, although there was a trend toward significance (*P=0*.*075*). Fish exposed to BIMU8 showed significant higher respiratory rate compared to fentanyl alone (*P=0*.*004*). The AMPA positive allosteric modulator CX614 did not significantly reverse respiratory rate depression by fentanyl (*P=0*.*073*, **Figure 4b**), but there was a trend toward significance. CX614/fentanyl showed lower respiratory rate compared to control (*P=0*.*002*), suggesting that CX614 at this dosage only partially reversed respiratory depression by fentanyl. CX614 alone presented higher respiratory rate when compared to fentanyl group (*P<0*.*001*). Higher dosages of CX614 induced seizure and were not tested. To determine whether the respiratory depression observed with fentanyl was due to a direct effect of fentanyl on respiratory rate rather than an indirect effect on sedation (Montandon & Horner, 2019) or pain circuits (Jiang, Alheid, Calandriello, & McCrimmon, 2004), we administered lidocaine, an analgesic not acting on MORs (Lopez-Luna et al., 2017). Lidocaine did not significantly change respiratory rate compared to control or vehicle/ethanol (*P=0*.*346*, **Figure 4c**).

**Figure 4.**
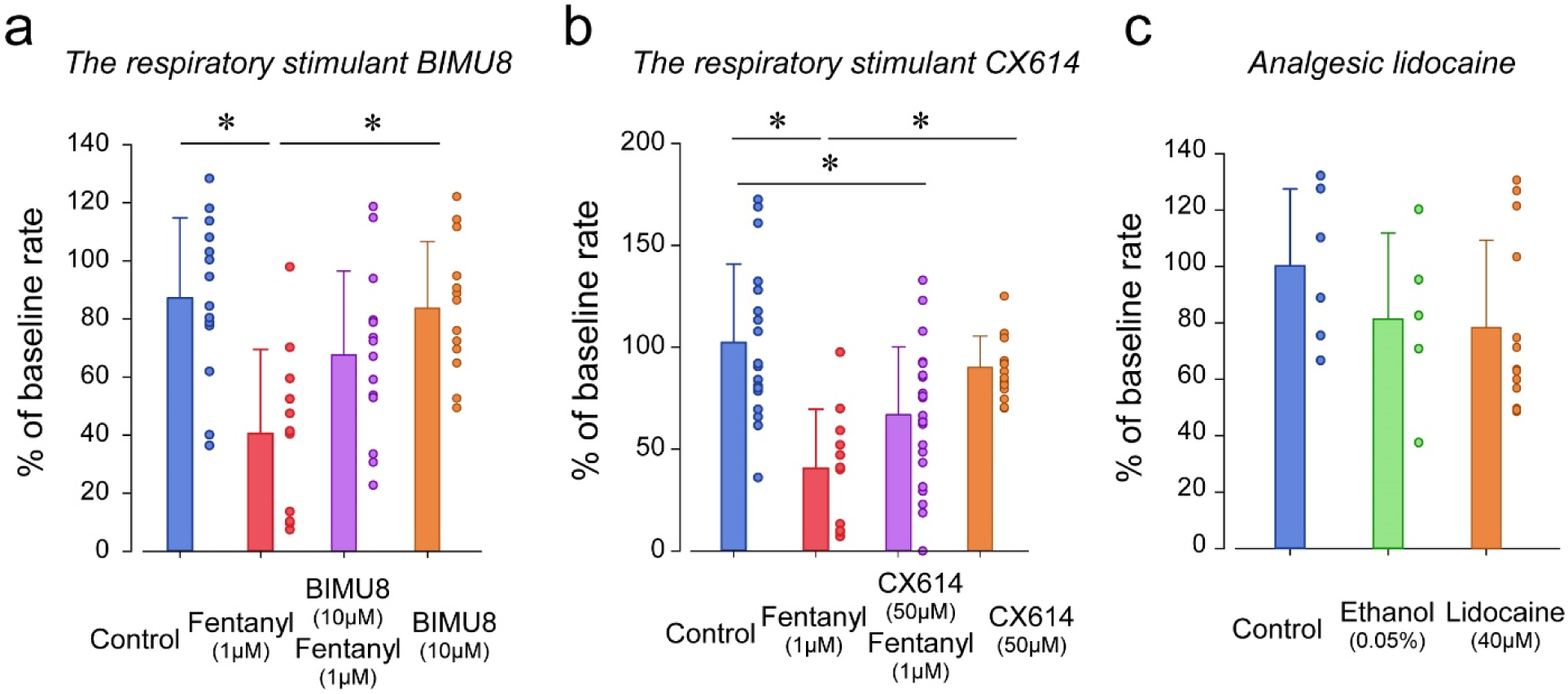
Respiratory depression and respiratory stimulants in larval zebrafish. The respiratory stimulant BIMU8 (10 µM), a 5-HT_4A_ serotonin receptor agonist, in combination with fentanyl (n=14) was compared to fentanyl alone (n=11) or BIMU8 alone (n=13) (**a**). The AMPA positive allosteric modulator CX614 (50 µM) and fentanyl (n=22) were also compared to fentanyl alone (n=11) or control (n=18, **b**). The analgesic lidocaine (n=12) was compared to ethanol (n=5) and control (n=6) (**c**). All data are presented as means ± standard deviation. Circles indicate individual data points for each zebrafish measured. * indicates groups significantly different with *P<0*.*05*.

To determine the analgesic properties of opioid drugs in larval zebrafish, we established a simple model of nociception combining exposure to formalin as a nociceptive stimulus and measurements of the subsequent swimming escape response. Using video recordings of larvae positioned in a multi-well plate, we tracked the individual movements of larvae exposed to embryo medium, formalin and a combination of formalin and fentanyl (**Figure 5**). We quantified two swimming behaviors (**Figure 5a**): swimming velocity (swimming distance in mm per second) and angular velocity (turn angle in degrees, per second). In the formalin group, larvae presented faster swimming velocity than control larvae (**Figure 5b**). Fentanyl combined with formalin showed considerable reduction in velocity compared to formalin alone (**Figure 5b**). Following exposure to formalin, swimming velocity significantly increased during the initial 3 minutes (**Figure 5c**, shaded blue), a response not observed in control larvae or formalin/fentanyl larvae (**Figure 5c**). We then compared the swimming velocity during the first 3 minutes following drug exposure for the different groups of larvae. The formalin group presented significantly faster velocity compared to the control group (*P<0*.*001*, **Figure 5d**). Velocity in the formalin/fentanyl group was significantly slower than when formalin was administered alone (*P=0*.*022*), suggesting that fentanyl reduced the escape response to formalin. Fentanyl administered alone did not decrease velocity when compared to control (*P=1*.*000*), showing that it did not affect swimming by itself. We then determined the effects of the µ-opioid receptor antagonists naloxone and CTAP (**Figure 5d**). The naloxone/formalin/fentanyl group presented significantly faster velocity than the control group (P<0.001) and was not significantly different than formalin alone suggesting that it blocked the effects of fentanyl. In CTAP/formalin/fentanyl group, there was significant differences compared to formalin/fentanyl or formalin alone (*P=1*.*000* and *P=0*.*069*).

**Figure 5.**
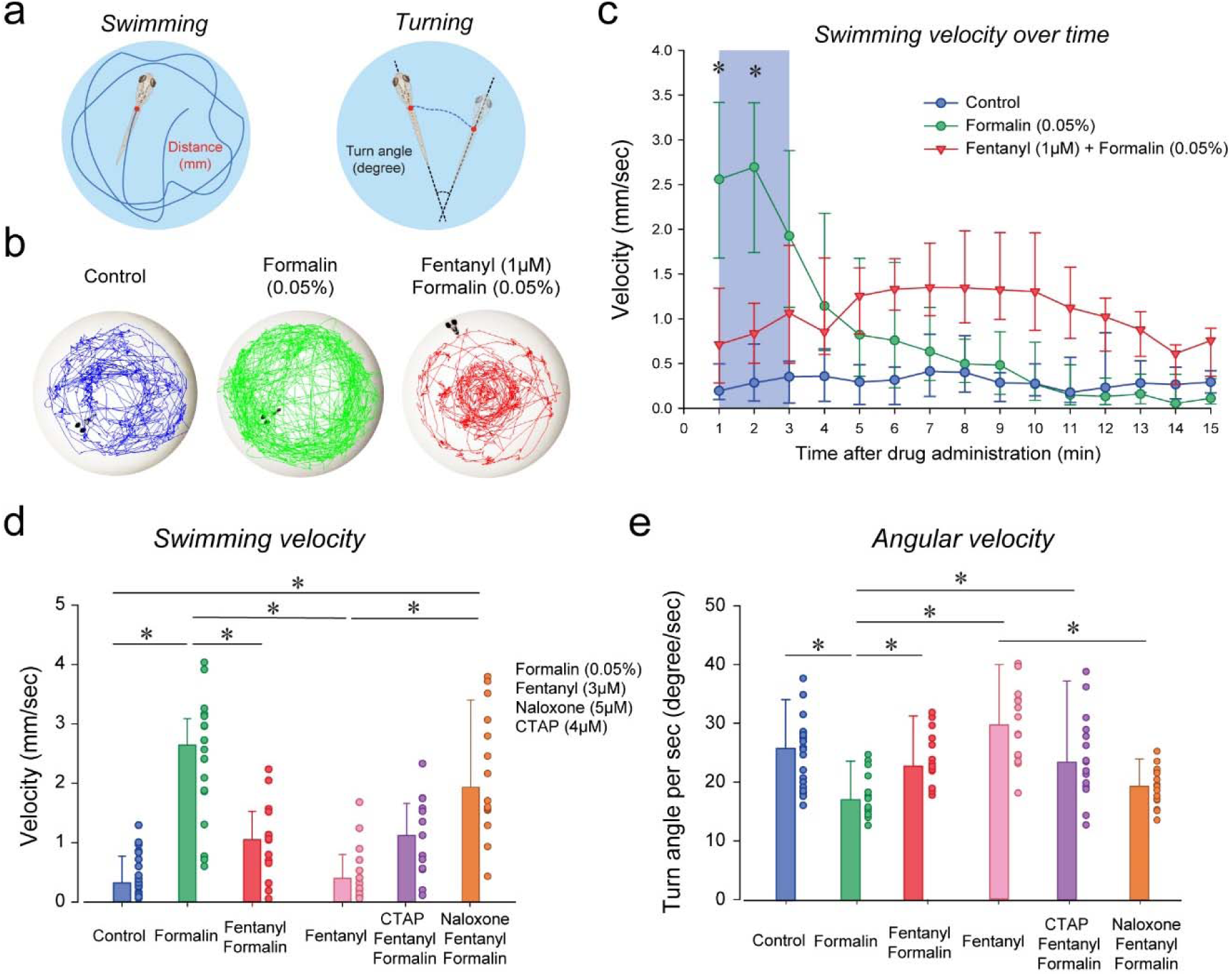
Analgesia by fentanyl in larval zebrafish. As an index of nociception in larval zebrafish, we measured behavioral responses to the nociceptive stimulus formalin. We quantified swimming and angular velocity in freely-moving larvae (**a**). Swimming velocity was quantified as the distance swam per second and angular velocity as the angle change of the fish every second. Control fish swam at a velocity ranging from 0.1 to 1.3 mm/sec, whereas swimming velocity was increased to 1.6-4.1 mm/sec with formalin. Addition of fentanyl reduced swimming velocity to 0.1-2.3 mm/sec (**b**). Following exposure to embryo medium only, formalin or formalin/fentanyl, swimming velocity was strongly increased by formalin within 3 minutes and was only moderately increased by fentanyl and formalin compared to controls (**c**). Comparison of averaged velocities for the first 3 minutes after baseline (shaded blue) showed that formalin (n=15) significantly increased velocity compared to the control (n=10), whereas fentanyl (n=12) reduced the formalin response (**d**). Naloxone blocked the effect of fentanyl (n=13), whereas CTAP (n=12) failed to block them. Angular velocity was significantly reduced by formalin an effect reversed by fentanyl/formalin or fentanyl alone (**e**). Naloxone blocked the fentanyl effects, but not CTAP. All data are presented as medians with error bars showing interquartile range. Circles indicate individual data points for each zebrafish measured. * indicates groups significantly different with *P<0*.*05*.

We then looked at the effect of formalin on angular velocity. The formalin group showed reduced turning velocity compared to control (*P<0*.*001*, **Figure 5e**), an effect that was reversed by fentanyl (*P=0*.*012*). Fentanyl administered alone had no effect of angular velocity (*P=1*.*000*). The naloxone/fentanyl/formalin group was not significantly different compred to the formalin group (*P=1*.*000*), suggesting that it blocked the effect of fentanyl. The CTAP/formalin/fentanyl group showed an increased turning velocity compared to formalin (*P=0*.*025*), suggesting that it failed to block the effect of fentanyl.

To determine whether larval zebrafish are sensitive to other types of analgesics, we exposed larvae to lidocaine, a non-opioid anesthetic (**Figure 6 a, b**). As previously shown, formalin increased velocity compared to control (*P<0*.*001*, **Figure 6b**). Lidocaine and formalin did not significantly change swimming velocity compared to formalin alone, although we observed a reduction in variability in the lidocaine/formalin group (*P=1*.*000*). Lidocaine by itself was not different from control (*P=1*.*000*). BIMU8/formalin/fentanyl did not significantly change swimming velocity compared to the control (*P=0*.*074*) or formalin/fentanyl group(*P=1*.*000*, **Figure 6c**). However, CX614/formalin/fentanyl group presented a higher swimming velocity than the control group (*P<0*.*001*) but did not show significant difference compared to formalin alone (*P=1*.*000*) or formalin/fentanyl (*P=0*.*168*). In Tübingen fish, formalin did not significantly increase swimming velocity (*P=0*.*151*, **Figure 6d**), but the combination of fentanyl and formalin did (*P=0*.*026*).

**Figure 6.**
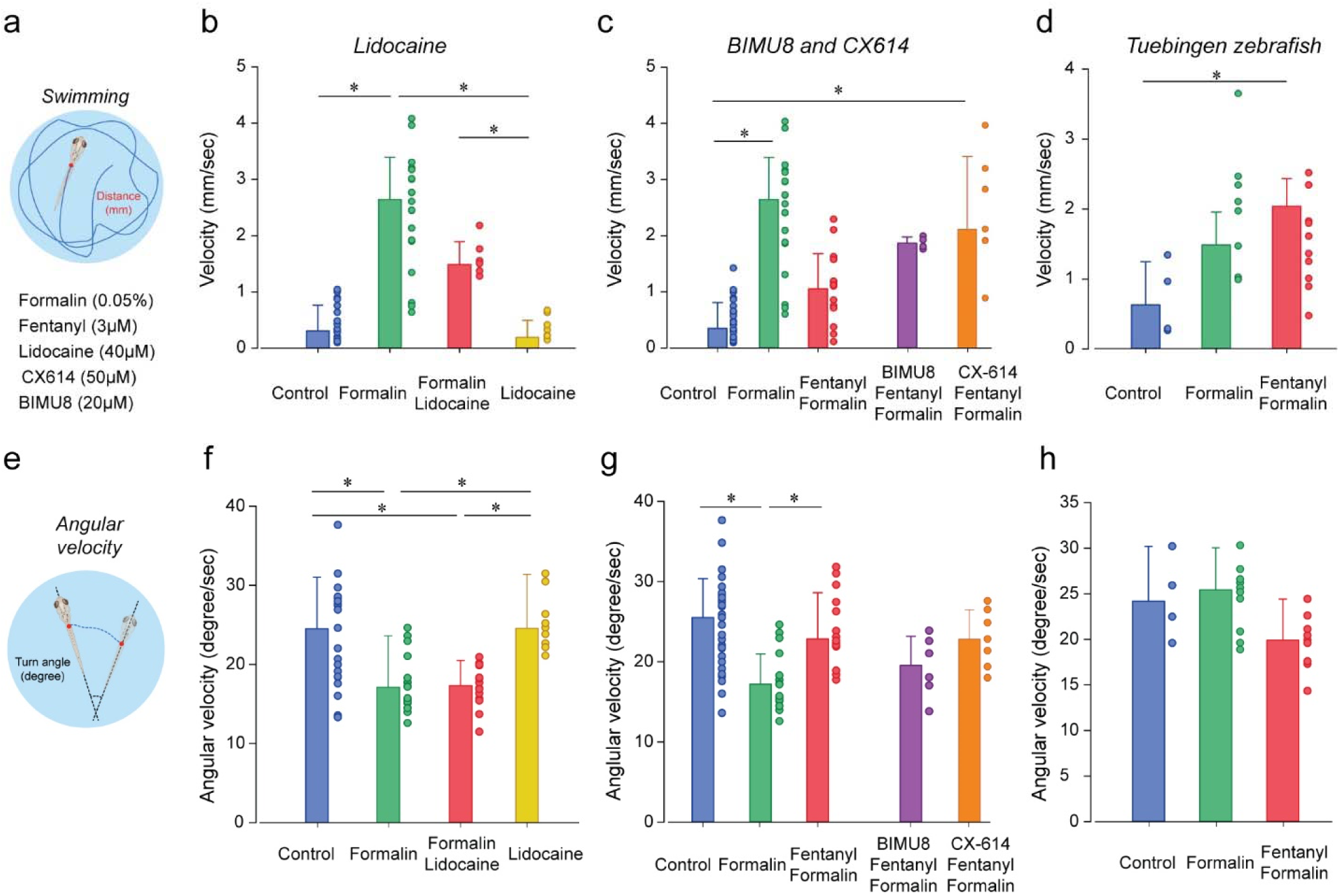
Analgesic profiles of larval zebrafish. To determine whether our analgesia assay can be confirmed with non-opioid analgesics, we used lidocaine, an anesthetic widely used in the clinic (**a**). Lidocaine only moderately reduced swimming velocity when administered with formalin (n=6, **b**). The respiratory stimulant BIMU8 (n=5) when administered with fentanyl and formalin did not significantly affect swimming velocity when compared to formalin alone (n=17). On the other hand, CX614 (n=7) presented significant differences compared to the control (n=24) (**c**). We also looked the analgesic properties of fentanyl in Tübingen zebrafish (**d**). As observed with respiratory assays, fentanyl (n=10) did not reduce the swimming response to formalin (n=7) as it did with AB zebrafish (d). All data are presented as medians with error bars showing interquartile range. Circles indicate individual data points for each zebrafish measured. * indicates groups significantly different with *P<0*.*05*.

We then assessed the effects of lidocaine on turning velocity (**Figure 6e, f**). Formalin decreased turning velocity (*P=0*.*005*), an effect not reversed by lidocaine/formalin (*P=1*.*000*). Turning velocity with lidocaine alone was not significantly different from the control (*P=1*.*000*). The stimulant BIMU8/formalin/fentanyl did not show any differences in turning velocity compared to formalin/fentanyl (P=0.238, **Figure 6g**) or formalin alone (*P=0*.*776*). Turning velocity with CX614 was also not significant different compared to formalin/fentanyl (*P=0*.*840*) or formalin alone (*P=0*.*086*). In Tübingen fish, turning velocity in formalin was different from fentanyl/formalin (*P=0*.*024*, **Figure 6h**).

## Discussion

Opioid drugs are widely used as analgesics but present the severe side-effect of respiratory depression that can be lethal with overdose. No safe opioid therapy currently exist to treat severe and chronic pain. To identify new potent pain treatments without side-effects, opioid drug discovery needs to explore new research avenues. Novel approaches to drug discovery have been recently used including structure-based drug discovery using computer simulations (Manglik et al., 2016) or cell assays (Winpenny, Clark, & Cawkill, 2016). Most drug discovery approaches use target-based approaches which exclude many drug candidates that do not specifically target proteins of interest. Here we propose phenotype-based drug discovery approaches in larval zebrafish to identify potent opioid pain therapies without the side-effects of respiratory depression. The larval zebrafish has emerged as a powerful model system for drug and gene discovery since it combines the biological complexity of *in vivo* models with a nervous system homolog to humans including the respiratory neural network. Such approach has the potential to validate many drug candidates without making any assumptions on their mechanisms of action or targets. Our drug discovery approach proposes to test whether drugs or combinations of drugs can be used as analgesics without the side-effects of respiratory depression and such strategy is of considerable interest (Dahan et al., 2018; (Montandon & Slutsky, 2019). We combined several assays to test the respiratory depressant effects of pain killers and their analgesics properties. We demonstrated that larval zebrafish (days post-fertilization 12-14) can be used to test the analgesic properties of opioid drugs and to test the severity of respiratory depression. Our assays provide simple, yet effective, ways to quickly test potential drug candidates in a complex animal model while preserving its central nervous system and brain blood barrier. Due to the high production of fish embryos, we can use these phenotypic assays to perform high-throughput drug screening protocols that are otherwise not feasible in other animal models. Importantly, drug screening in rodents is not feasible as it would require extensive resources and time. Here, we propose new behavioral assays in larval zebrafish where live animals with intact central nervous system can be leveraged for phenotype-based drug discovery combining respiratory depression and analgesia by opioid drugs.

Respiratory depression by opioids has been widely studied in mice, rats, dogs and sheep (Krause et al., 2009; (Montandon et al., 2011; (Montandon, Ren, et al., 2016). The use of complex animal models to assess respiratory variables comes with many challenges. To accurately assess respiratory depression by opioid drugs, it is critical to quantify respiratory variables without the use of anesthetics which interact with the respiratory system, drug metabolism, and mechanisms of action of opioid drugs. Respiratory assessments in non-anesthetized rodents is the gold standard but it raises concerns due to the stress associated with animal handling, opioid injection, and changes due to altered arousal states and behaviours (Montandon, Cushing, et al., 2016; (Montandon & Horner, 2019), which may ultimately affect research outcomes. In larval zebrafish, we demonstrated, for the first time, that respiratory network activity can be assessed, that larval zebrafish present respiratory rate depression by opioid drugs in a similar fashion that humans, and that opioid pharmacology observed in humans is preserved in zebrafish. Most fishes have a complex respiratory system to move water through their gills. Although fishes use a different strategy to absorb oxygen and eliminate carbon dioxide than mammals, they rhythmically produce mandibular movements to move water through their gills. In lampreys, the paratrigeminal respiratory group (pTRG) generates rhythmic mandible movements (Bongianni et al., 2016). Because of their close evolutionary origins (Missaghi et al., 2016), the pTRG shares similarities with the mammalian respiratory network (Cinelli et al., 2013) which generates breathing and regulates respiratory depression by opioids (Montandon et al., 2011). The pTRG presents similar functional properties than mammals (Gray, Rekling, Bocchiaro, & Feldman, 1999) such as sensitivity to substance P (Mutolo et al., 2010). Here, we propose that respiratory mandible movements controlled, at least in great part, by the pTRG can be used as an index of respiratory network activity, similarly to respiratory activity of the trigeminal muscle in mammals (Jacquin et al., 1999). We showed that the rate of respiratory movements presented dose-dependent decreases in response to increasing dosages of the MOR ligand fentanyl. Although it is difficult to compare administration of fentanyl through fish water in zebrafish and systemic injection in mammals, the dosages used in zebrafish corresponds to those used in rodents (Yassen, Olofsen, Kan, Dahan, & Danhof, 2008). Respiratory depression by fentanyl was substantially blocked by the MOR antagonists CTAP and naloxone, which is consistent with the fact that fentanyl is acting mostly through MORs (James & Williams, 2020). Interestingly, morphine did not reduce significantly respiratory rate in larval zebrafish at the dosage used in this study, which is consistent with the less potent properties of morphine, but not with the mild respiratory depression usually observed with morphine in humans (Montandon, Cushing, et al., 2016). Fentanyl presents high affinity for MOR and this receptor is found in larval zebrafish (Gonzalez-Nunez, Jimenez Gonzalez, Barreto-Valer, & Rodriguez, 2013). Morphine has a lower affinity to MOR than fentanyl which may explain the lack of respiratory depressant effect of morphine in zebrafish.

The two major strains of wild-type zebrafish, AB and Tübinger (TU), showed different sensitivities to fentanyl. AB fish present pronounced respiratory depression, an effect not observed in TU fish, regardless of the concentration used (data not shown). This strain-specificity suggests that the mechanisms of action of opioid drugs differ between zebrafish strains which could be due to polymorphisms of the MOR gene as it can be found in humans (Oertel, Schmidt, Schneider, Geisslinger, & Lotsch, 2006) or genetic differences in various genes involved in MOR inhibition (Bian, Wu, Su, & Li, 2012). To determine whether respiratory stimulants can reverse respiratory depression by opioids in larval zebrafish, we tested two types of agonists targeting excitatory receptors: ampakines which comprise drugs acting on AMPA receptors (Ren, Poon, Tang, Funk, & Greer, 2006) and serotoninergic agents (Manzke et al., 2003). CX-614, an ampakine allosteric modulator of AMPA receptors, did not reverse respiratory depression in our larval zebrafish models, which is not consistent with its effects in rodents (Ren et al., 2006) or humans (Dahan et al., 2018). This lack of ampakine effects may be due to the low concentration used in this study because higher doses of CX-614 were inducing severe seizures in larval zebrafish (data not shown). BIMU8, a 5-HT_4A_ agonist, was administered in combination with fentanyl and compared with fentanyl alone. BIMU8 did not significantly reverse respiratory depression at the concentrations tested, which is consistent with the low efficacy of this treatment in humans (Lotsch, Skarke, Schneider, Hummel, & Geisslinger, 2005). Other serotonin agents such as 5-HT_1_ and 5-HT_3_ agonists (van der Schier, Roozekrans, van Velzen, Dahan, & Niesters, 2014) can be easily tested using our zebrafish models.

Zebrafish larvae were previously used as animal models to study pain and analgesia. Acetic or citric acids were added to fish water to induce pain (Lopez-Luna et al., 2017; (Steenbergen & Bardine, 2014). Our models did not use acids as nociceptive stimuli as they change water pH, to which zebrafish are sensitive (Avdesh et al., 2012; (Steenbergen & Bardine, 2014). Instead, we used formalin, a chemical widely used to induce pain in rodent models (Yoon et al., 2015), and showed that formalin increased swimming velocity. Here, we suggest that increased swimming velocity in response to formalin represents the fish escape response to nociceptive stimuli. Fentanyl reduced the swimming response to formalin suggesting that it reduced pain induced by formalin. MOR antagonists abolished the inhibitory effects of fentanyl on the swimming response to formalin demonstrating that it may block the analgesic effects of fentanyl. These results are consistent with the analgesic effects of fentanyl and the blocking effects of naloxone observed in humans (Gutstein, 2001). Formalin is however not an ideal nociceptive stimulus as it may induce effects not related to nociception. It could be suggested that the escape response observed in our assays is due to the effect of formalin on locomotor activity (not nociception) and that the reduced response with fentanyl may be due to inhibition of motor activity. However, fentanyl alone did not reduce swimming velocity compared to control larvae, suggesting that the effect of fentanyl on the formalin response is not due to reduced locomotion. Lidocaine, a non-opioid analgesic, applied with formalin did not show a significant reduction compared to formalin alone. Lidocaine induces analgesia by inactivating voltage-gated sodium channels in neurons (Tetzlaff, 2000), and it is unclear whether sodium channels are involved in pain circuits in larval zebrafish. As observed with the respiratory depression assays, the two stimulants BIMU-8 and CX-614 did not reverse analgesia which is consistent with their effects in rodents (Dahan et al., 2018). BIMU-8 administered alone produced a slight increase in swimming velocity when compared with control conditions, which is consistent with the role of serotoninergic agents on locomotion (Perrier & Cotel, 2015). In conclusion, we established opioid analgesia assays that can be used to quantify pain and the analgesic properties of new opioid therapies that can easily be used for large scale drug discovery using larval zebrafish.

Opioid analgesics constitute essential pain therapies that present the lethal side-effect of respiratory depression therefore limiting their effective use in the clinical and at-home settings. There is currently no effective safe pain therapy due to the difficulty at identifying new drug combinations with potent analgesia but reduced respiratory side-effects (Dahan et al., 2018). Using larval zebrafish, we propose models allowing phenotype-based drug discovery approaches, i.e. permitting drug testing without assumptions related to mechanisms of action and targets. Our novel drug discovery models allow high-throughput drug screening in a simple and amenable animal model presenting similar pharmacological and genetic profiles than humans (MacRae & Peterson, 2015). Although zebrafish has been used to identify new anesthetics and their mechanisms of actions (McGrath et al., 2020; (Yang et al., 2019), our study is the first to assess respiratory depression by opioid drugs in larval zebrafish combined with analgesia. In addition to high-throughput screening approaches, our models can also be used in combination of transgenic or knockout zebrafish to better understand the mechanisms opioid inhibition, analgesia and respiratory depression, as well as live microscopy of neural circuits of pain and respiration (Ahrens, Orger, Robson, Li, & Keller, 2013).

## Acknowledgements

Research was supported by the Research Innovation Council Fund of St. Michael’s Hospital Foundation and the J. P. Bickell Foundation Medical Grant. SZ was supported by a Ontario Graduate Scholarship. We thank Alexandra Ho for her help with data analysis.

